# Substrate elasticity regulates cytoskeletal remodeling and mechanical behavior of U2OS osteosarcoma cells

**DOI:** 10.1101/2025.09.22.677759

**Authors:** Carina Rząca, Agata Kubisiak, Dominik Panek, Marta Targosz-Korecka, Zenon Rajfur

## Abstract

Substrate elasticity plays a pivotal role in regulating the morphology, mechanical properties, and cytoskeletal organization of cancer cells. In this study, we examined the response of U2OS osteosarcoma cells to substrates of varying stiffness, with a particular focus on cytoskeletal remodeling, cell elasticity, and microparticle internalization. To simulate environments of moderate and high stiffness, cells were cultured on polyacrylamide (PA) hydrogels with a stiffness of 40 kPa and on rigid glass substrates, respectively.

Changes in cell morphology and cytoskeletal organization were assessed using fluorescence microscopy, while cell mechanical properties were measured using atomic force microscopy (AFM). To investigate the relationship between substrate mechanics and endocytic activity, carboxylated fluorescent 2 µm latex microspheres were introduced to the cell culture system.

Our study showed that cell spreading increased with substrate stiffness. U2OS cells cultured on glass exhibited a significantly larger surface area, more actin stress fibers, and a more organized, stretched cytoskeletal architecture compared to cells grown on 40 kPa PA gels. AFM measurements further demonstrated that cells on glass were mechanically stiffer than those on PA substrates. Microparticle uptake was also strongly influenced by substrate stiffness. Cells cultured in the 40 kPa PA gels internalized a significantly greater number of fluorescent microspheres. Notably, these cells frequently formed distinct, cup-like structures composed of microtubules around the beads. Three-dimensional image reconstructions revealed that these structures encapsulate the particles in an asymmetrical manner, indicative of active cytoskeletal remodeling.

To better understand the molecular composition of these microtubule-based structures, we analyzed the localization of selected microtubule-associated proteins (MAPs), including IQGAP1, CLIP1, and MARK2 (Conboy J.P. et al. 2024). Interestingly, only IQGAP1 was localized prominently to the microtubule cups on 40 kPa gels, often forming ring-like structures surrounding the beads. In some cases, these rings were observed independently of detectable microtubules, suggesting the involvement of an active, possibly microtubule-initiated, endocytic process.

In conclusion, our findings demonstrate that substrate stiffness modulates multiple aspects of U2OS cell behavior, including morphology, cytoskeletal arrangement, mechanical properties, and microparticle uptake. These results underscore the mechanosensitive nature of osteosarcoma cells and highlight novel roles for microtubule structures and MAPs, particularly IQGAP1 in stiffness-dependent cellular uptake mechanisms.

## 1. Introduction

Cells have an innate ability to detect and respond to physical and chemical signals from their surroundings. Through processes such as mechanosensing and mechanotransduction, cells can sense the mechanical properties of their environment and convert these signals into biochemical reactions. Despite continuous progress in understanding of how cells interact with mechanical cues, the mechanisms by which they perceive changes in matrix stiffness are still being explored. Recent research has focused on using substrate gels with adjustable elasticity to replicate the mechanical properties of native tissues, shedding light on the impact of substrate stiffness on cellular behavior (Venugopal et al., 2018).

The elasticity of tissues in mammals varies significantly, which is reflected in their Young’s modulus (E) values. For example, the brain has a Young’s modulus of 0.1–1 kPa (Barnes et al., 2017; Budday et al., 2015), muscles are 20-40 kPa (Basford et al., 2002; Horikawa et al., 1993), lungs show values between 1-5 kPa (Yu et al., 2017) and bones exhibit much higher stiffness at 14–23 GPa (Shoaib et al., 2022). Tumor cells, such as osteosarcoma (OS), are of particular interest due to their ability to metastasize to various organs, encountering a wide range of tissue stiffness. Understanding how these variations in stiffness influence cellular behavior is key to advancing knowledge of mechanotransduction and improving therapeutic strategies for tissue repair and cancer treatment (Shoaib et al., 2022).

The cytoskeleton, which consists of actin filaments, microtubules, and intermediate filaments, plays a crucial role in maintaining cell structure and function by transmitting mechanical forces. Substrate stiffness can greatly affect cell behavior, influencing processes such as differentiation, disease progression, and tissue regeneration (Fletcher & Mullins, 2010). For successful tissue engineering and regenerative therapies, it is essential to replicate mechanical cues that match those found in native tissues. While much is known about how chemical signals influence cellular functions, the impact of mechanical cues from the extracellular matrix on cell behavior is less well understood. Recent studies have demonstrated that cells can adjust their cytoskeletal organization and mechanical properties in response to changes in the stiffness of their environment (Shou et al., 2023).

Latex beads are commonly used in biomedical research. These beads, with a diameter typically in the micrometer range, are especially valuable in studies related to phagocytosis and mechanobiology. These microspheres serve as model objects to investigate cellular engulfment processes and enable the controlled study of endocytose mechanisms. A well-established technique involving microspheres is tether extraction, in which a bead is attached to a cell membrane, and a membrane-derived thread is pulled from the bead to study membrane viscosity and actin–membrane interactions (Gruenberg & Maxfield, 1995). In examining cytoskeletal rearrangements, beads are frequently used to observe how they influence cellular structures.

It is well established that actin can form phagocytic cups, particularly in macrophage cells. However, a previous study by Adamczyk et al. (2021) reported the formation of cup-like structures composed of microtubules in fibroblast cells following interaction with polystyrene microspheres. In the present study, we identified a similar phenomenon in U2OS osteosarcoma cells. The primary objective of our research was to investigate how microspheres influence the cellular architecture of cancer cells, with a particular focus on microtubules and microtubule-associated proteins (MAPs). Additionally, we examined the role of substrate stiffness in modulating these interactions. Notably, we observed the formation of microtubule-based cup-like structures—an organization not typically associated with endocytosis—with a higher frequency on elastic polyacrylamide substrates compared to rigid glass. MAPs such as MARK2, CLIP1, and IQGAP1 proteins are known regulators of microtubule dynamics and cytoskeletal organization. In our experiments, we observed accumulation of IQGAP1 around polystyrene microspheres, even in the absence of microtubules, suggesting a microtubule-independent role for this protein in vesicle or particle recognition. These findings point to the novel role of both microtubules and MAPs in mechanotransduction and highlight their involvement in cytoskeletal remodeling in response to substrate stiffness.

Importantly, the novelty of our work extends beyond morphological characterization by integrating atomic force microscopy (AFM) to correlate phenotypic observations with intracellular mechanical properties. This integrative, multidimensional approach offers new mechanistic insights into how U2OS osteosarcoma cells perceive and adapt to mechanical cues in their environment, establishing a robust model for investigating mechanotransduction processes relevant to cancer progression.

## 2. Materials and methods

### 2.1. Cell culture

The osteosarcoma (U2OS) cell line was obtained from ATCC. Cells were cultured in McCoy’s 5a Medium (L0066, Biowest) supplemented with 10% Gibco fetal bovine serum (FBS) (10270106, Thermo Fisher Scientific), and 100 I.U./mL Penicillin and 100 μg/mL Streptomycin Solution (L0022, Biowest) at 37 °C, 5% CO_2_, and humidity in an incubator.

### 2.2. Polyacrylamide Gel Substrate Preparation

Glass-bottomed dishes were treated with a solution of 3-(trimethoxy silyl) propyl methacrylate (440159, Sigma-Aldrich), 99.5% acetic acid (010330, Sigma-Aldrich) and 96% ethanol (64175, Sigma-Aldrich) in a ratio of 1:1:14 for 30 minutes, washed twice with 96% ethanol and dried. Polyacrylamide substrates were prepared by mixing acrylamide, bis-acrylamide, HEPES, TEMED and APS, and the proportions of these components for the preparation of 40 kPa substrate was taken from the protocol (Tse & Engler, 2010). In the next step, 22 µL of the solution was placed in the centre of a 35 mm glass bottom dishes with 20 mm well, and a thickness of #0, designed for high-resolution imaging (Cellvis, D35-20-0-N) and then covered with a glass coverslip and incubated for 1 hour at room temperature. Next, the coverslip was removed, and the gel was incubated with 2 µg/mL Sulfo-SANPAH (22589, Thermo Fisher Scientific) under 365 nm ultraviolet (UV) irradiation for 5 min. After that, PA gels were washed with sterile HEPES and phosphate-buffered saline (PBS) solution 3 times, next incubated with 10 µg/mL fibronectin at room temperature for 2 h. Cells were then seeded on 35 mm glass-bottom dishes at the concentration of 2 * 10^4^ cells onto gel for 0 h, 6 h, 12 h, 18 h and 24h. Elasticity of the gel was confirmed using AFM, substrates Young’s Modulus was 38.84 ± 5.59 kPa (Supplementary file 1). As these values fall within the margin of error, the designation of 40 kPa has been retained.

### 2.3. Microspheres

Fluoresbrite YG Carboxylate Microspheres with a diameter of 2 μm were obtained from Polysciences (09847, Polysciences Inc.). Fluoresbrite® carboxylate microspheres are fluorescent monodispersed polystyrene microspheres that have carboxylate groups on their surfaces which can be activated for the covalent coupling of proteins. These microspheres suspension concentration was 5.68 x 10^9^ particles/ml. Excitation max. = 441nm and emission max. = 486nm. Microparticles were added to cells between 12h and 18h post-seeding and incubated for 3h. After this time cells were fixed using 4% formaldehyde solution.

### 2.4. Cell Labeling and Immunofluorescence

Cells were fixed with 4% formaldehyde solution (47608, Sigma-Aldrich) for 20 minutes at 37°C in an incubator and subsequently permeabilized with 0.1% Triton X-100 (T8787, Sigma-Aldrich) for 5 minutes to enable intracellular access for antibodies and other labeling reagents. To block non-specific binding sites, cells were incubated in 4% BSA in PBST buffer (PBS with Tween-20; A7906, Sigma-Aldrich) for 1 hour at room temperature (RT).

Primary antibody staining was performed overnight at 4°C. Cells were incubated with the following primary antibodies diluted in 3% BSA in PBS: anti-α-tubulin (1:500; T9026, Sigma-Aldrich), anti-IQGAP1 (1:100; SAB4200079, Sigma-Aldrich), anti-MARK2 (1:100; SAB4500761, Sigma-Aldrich), and anti-CLIP1 (1:100; HPA026678, Sigma-Aldrich).

Following primary antibody incubation, cells were washed and incubated for 1 hour at RT in the dark with fluorophore-conjugated secondary antibodies: Alexa Fluor 555-labeled goat anti-mouse IgG (H+L) (1:200; A21422, Thermo Fisher Scientific) and Alexa Fluor 633-labeled goat anti-rabbit IgG (H+L) (1:100; A21070, Thermo Fisher Scientific).

For actin visualization, cells were incubated with DyLight 633-conjugated phalloidin (1:200; 21840, Thermo Fisher Scientific) for 1 hour at RT.

### 2.5. Atomic Force Microscopy

The measurements were carried out using fixed cells in Hanks’ Balanced Salt Solution (H8264, Sigma-Aldrich) as a measuring medium, according to the procedure described in the paper by (Targosz-Korecka et al., 2015). All experiments were done in at least 3 replicates. The measurement was carried out in a drop of medium with NanoWizard 3 NanoScience AFM (JPK Instruments). Cells and substrates imaging was performed using pyramidal-shaped Pt-Ir coated cantilevers (SCM-PIC-V2, Bruker) with a nominal spring constant of 0.1 N/m. Measurements were conducted using a force mapping mode. For each cell, a spatial map of force vs distance (FD) curves at grid of 64 × 64 points was measured. The size of the grid corresponds to a square surface with dimensions ranging from 30 μm × 30 μm and 40 μm × 40 μm. To calculate the cell’s elastic modulus E (reduced Young’s modulus), the set of force-distance curves was analyzed by fitting them to the Hertz-Sneddon model, as detailed in our earlier publications (Targosz-Korecka et al., 2017). The results were presented as specialized elastic modulus maps (a qualitative representation of the data) and as the average elastic modulus value calculated for each cell (a quantitative analysis). Consequently, for each measured cell, we obtained topographical images that correlated with the elasticity map. Force-distance curves were measured at a speed of 10 µm/s and the maximal applied force was 1.3 nN. Before a series of force-distance measurements, calibration of cantilever spring constant was performed by using Nanowizard software. The topography and elastic modulus images were generated using JPK Data Processing Software version 5.1 (https://jpkspm-data-processing.software.informer.com/). The position of scan areas was continuously controlled by inverted optical microscope 10x, 1.4 NA and 40x NA Olympus IX71 (Olympus, Japan).

### 2.6. Microscopy and Images Analysis

Confocal images were acquired using Zeiss LSM 710 confocal module set on Zeiss Axio Observer.Z1 inverted microscope (Carl Zeiss Microscopy GmbH, Germany) using an oil immersion 40x, 1.4 NA Plan-Apochromat objective. To measure the cell height, z-stacks were taken at an interval of 0.3 µm. The height was calculated as the distance between the top and bottom cell edge in ImageJ software.

Time-lapses of the live cells were taken using Zeiss Axio Observer Z1 (Carl Zeiss) with HXP 120 V and arc lamp (HXP R 120 W/45 C VIS, OSRAM), using an oil immersion 40x, 1.4 NA plan-apochromat objective and Hamamatsu ORCA-Flash 4.0 camera. For moving the stage and maintaining the focus, WSB PiezoDrive CAN and fc12 were used (Carl Zeiss), respectively. Heating Insert P Lab-Tek™ S (Pecon, Germany) together with TempModule S (Pecon, Germany) and CO_2_ Module S (Pecon, Germany) provided the right conditions for live cell imaging: 37 °C and 5% CO_2_.To analyze cell spreading area, firstly Cellpose software was used (Stringer et al., 2020) version 3.0, to obtain masked images. Acquired images were further analyzed using Python (version 3.11) and libraries such as numpy (version 1.26.4), scikit image (version 0.18.1), and OpenCV (version 4.5.5) and cell area was determined as the number of pixels in the image, where for each cell a unique numerical value was assigned. In such a way, counting the number of unique pixel values was equivalent to the area of a given cell.

### 2.7. Statistical Analysis

All tests were carried out in at least triplicate, and the results were averaged. Two-way ANOVA analysis followed by the Bonferroni post hoc test was used for comparisons. p values less than 0.05 were considered to be statistically significant. Data are presented as mean ± standard deviation (SD). All calculations and visualizations were done in Origin software (version 2023b).

## 3. Results

### 3.1. Substrate Stiffness Modulates Cell Morphology and Spreading

Brightfield time-lapse imaging revealed that U2OS cells progressively altered their morphology depending on substrate stiffness (Fig. 1A). Initially (0 h), cells displayed comparable areas across conditions: 194.76 ± 38.09 µm² (glass), and 247.36 ± 60.78 µm² (40 kPa PA). Already after 6 hours, U2OS cells exhibited a 3.5-fold increase in spreading area compared to their initial state, indicating that during this time cells were able to adhere, sense the substrate properties, and undergo flattening. Over the course of 24 hours, cell spreading further increased significantly, particularly on stiffer substrates such as glass (Fig. 1B). At 24 h, cells on glass exhibited the largest spreading area (1279.61 ± 380.09 µm²), and U2OS cells on 40 kPa PA (1069.58 ± 410.40 µm²) significantly less spread. Adherent cells typically require approximately 12 to 24 hours to firmly attach and initiate normal metabolic activity. It has been observed that cells reach a growth plateau around 24 hours post-seeding (Fig. 1).

**Figure 1.**
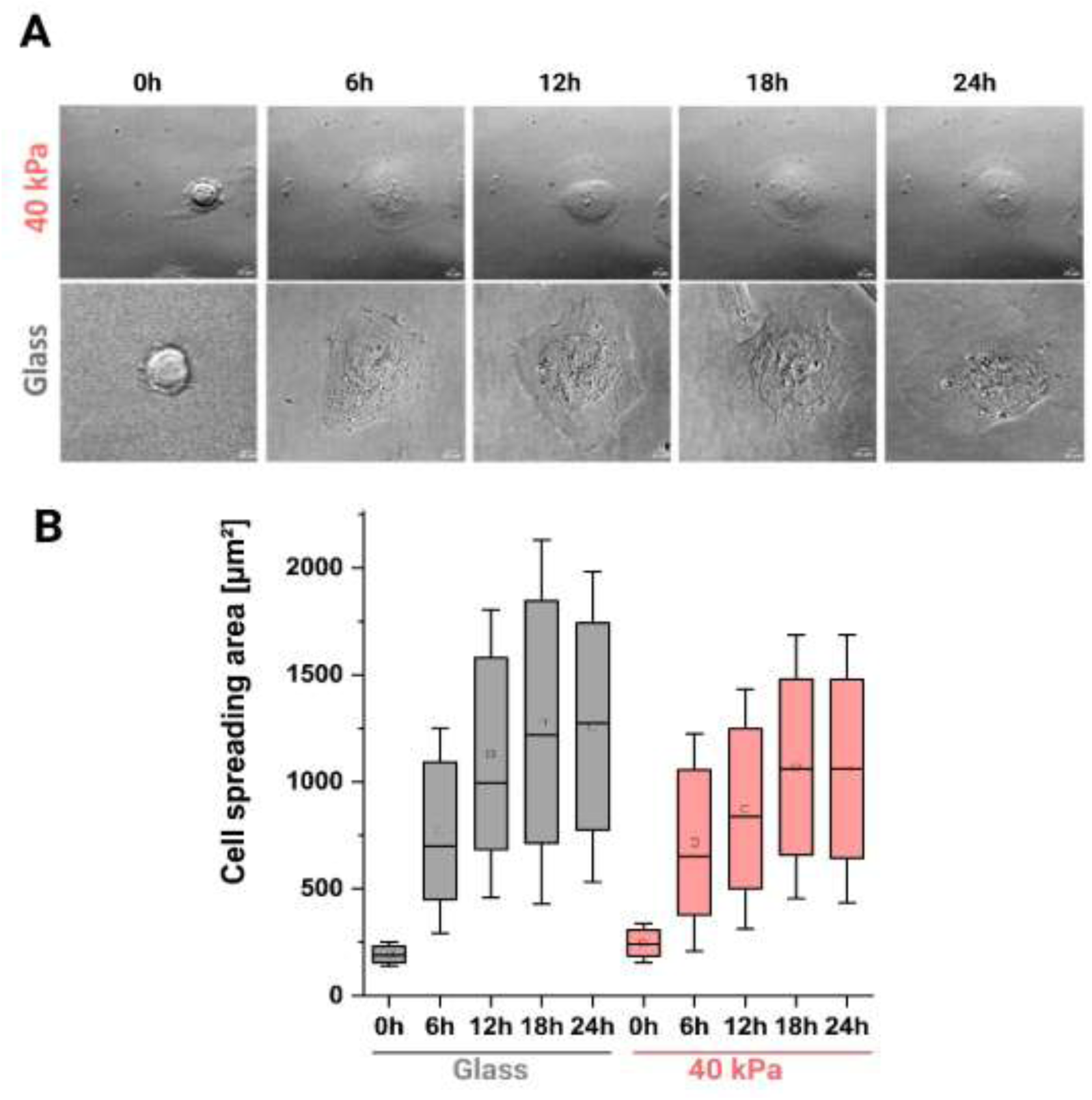
Effect of substrate stiffness on U2OS cell morphology and spreading area. (A) Representative images from time-lapses depicting changes in morphology and spreading behavior of U2OS cells on glass and 40 kPa PA gel substrates over 24 hours. Scale bars represent 20 μm. (B) Quantification of U2OS cell area on 40 kPa PA and glass substrates. Data are presented as mean values (square inside the boxes) with median values (line). Box represents the spread of 50% of the data (between 25–75 percentile values). Glass N=244, 40 kPa PA N=207. Statistical analysis was performed using two-way ANOVA followed by Bonferroni post hoc test (statistical significance is shown in Supplementary file).

### 3.2. Cytoskeletal Architecture Is Altered by Substrate Stiffness

Confocal fluorescence microscopy images (Fig. 2 A,B) demonstrated different arrangement of microtubules, vimentin, and actin filaments in cells at 12- and 24-hours post-seeding. At 12 hours post-seeding, confocal fluorescence microscopy images revealed that the arrangement of microtubules, vimentin, and actin filaments varied across substrates. On glass, cells displayed well-organized networks of all three cytoskeletal components, with dense actin structures and microtubule fibers extending from the nucleus throughout the cell body. On the 40 kPa PA substrate, the cytoskeletal networks were present but appeared slightly less organized than on glass, with a less prominent actin network. By 24 hours post-seeding, further differences in cytoskeletal organization became evident. Cells on glass retained their well-organized networks of microtubules, vimentin, and actin, with dense actin structures and microtubule fibers continuing to extend throughout the cell body.

**Figure 2.**
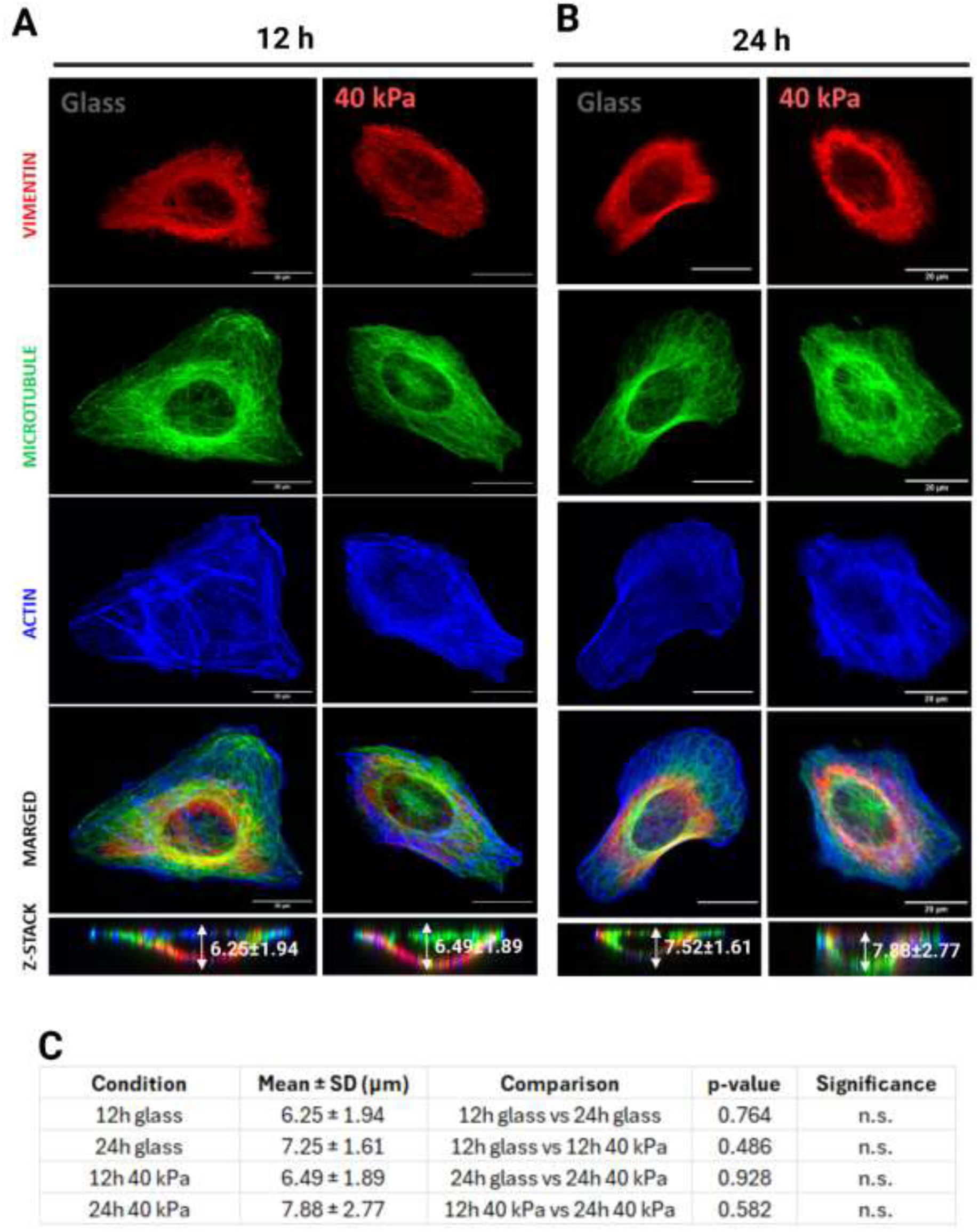
Cytoskeleton architecture of U2OS cells on different PA substrates. Representative images of the cytoskeleton of U2OS cells at 12 h (A) and 24 h (B) post-seeding are presented as a sum of z-stacks. The actin structure is represented in blue, the microtubules in green, and vimentin in red. Scale bars represent 20 μm. (C) Quantification of U2OS cell thickness [µm]. The statistics were calculated based on two-way ANOVA along with the post hoc test Bonferroni. n.s. – not significant (p > 0.05). Glass N=24, 40 kPa PA N=20.

Cross-sectional analysis demonstrated a consistent distribution of cytoskeletal components (Fig. 2C). At 12 hours post-seeding, cells cultured on the 40 kPa PA substrate exhibited a significantly greater mean z-stack height (7.88 ± 2.77 µm) compared to those on glass (6.25 ± 1.94 µm), indicating that cells on the softer substrate were less spread and exhibited increased height. After 24 hours, the mean cell height on glass slightly increased to 6.24 ± 1.89 µm, whereas it decreased on the 40 kPa substrate to 7.52 ± 1.61 µm. These findings suggest that substrate stiffness markedly influences cytoskeletal organization and cell morphology dynamics over time.

### 3.3. Substrate Stiffness Influences Cell Elasticity

AFM measurements (Fig. 3) revealed marked differences in the mechanical properties of U2OS cells depending on substrate stiffness. Cells cultured on glass exhibited the highest mean Young’s modulus values, indicating increased stiffness relative to those on compliant substrates (Fig. 3A). In contrast, cells on 40 kPa PA gels showed lower elastic modulus values, suggesting that cells actively respond to and adapt their mechanical state in accordance with substrate rigidity. Notably, even after 6 hours of culture, cells displayed substrate-dependent mechanical characteristics, with measured stiffness values of 835.55 ± 199.26 kPa on glass and 457 ± 206 kPa on 40 kPa PA gels. Over time (6 h, 12 h, and 18 h), U2OS cells exhibited oscillatory changes in elasticity across all tested substrates. However, the amplitude of these variations was more pronounced on glass than on 40 kPa PA gels, indicating more stable mechanical behavior on softer substrates.

**Figure 3.**
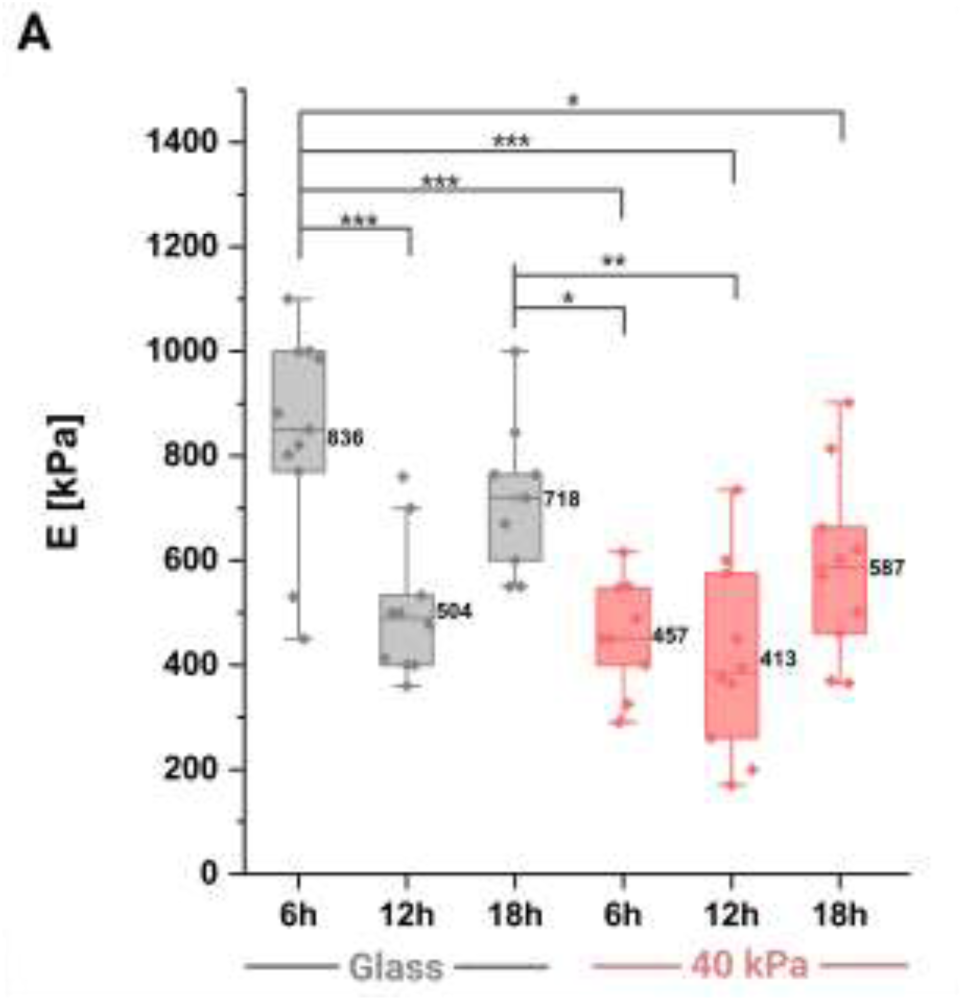
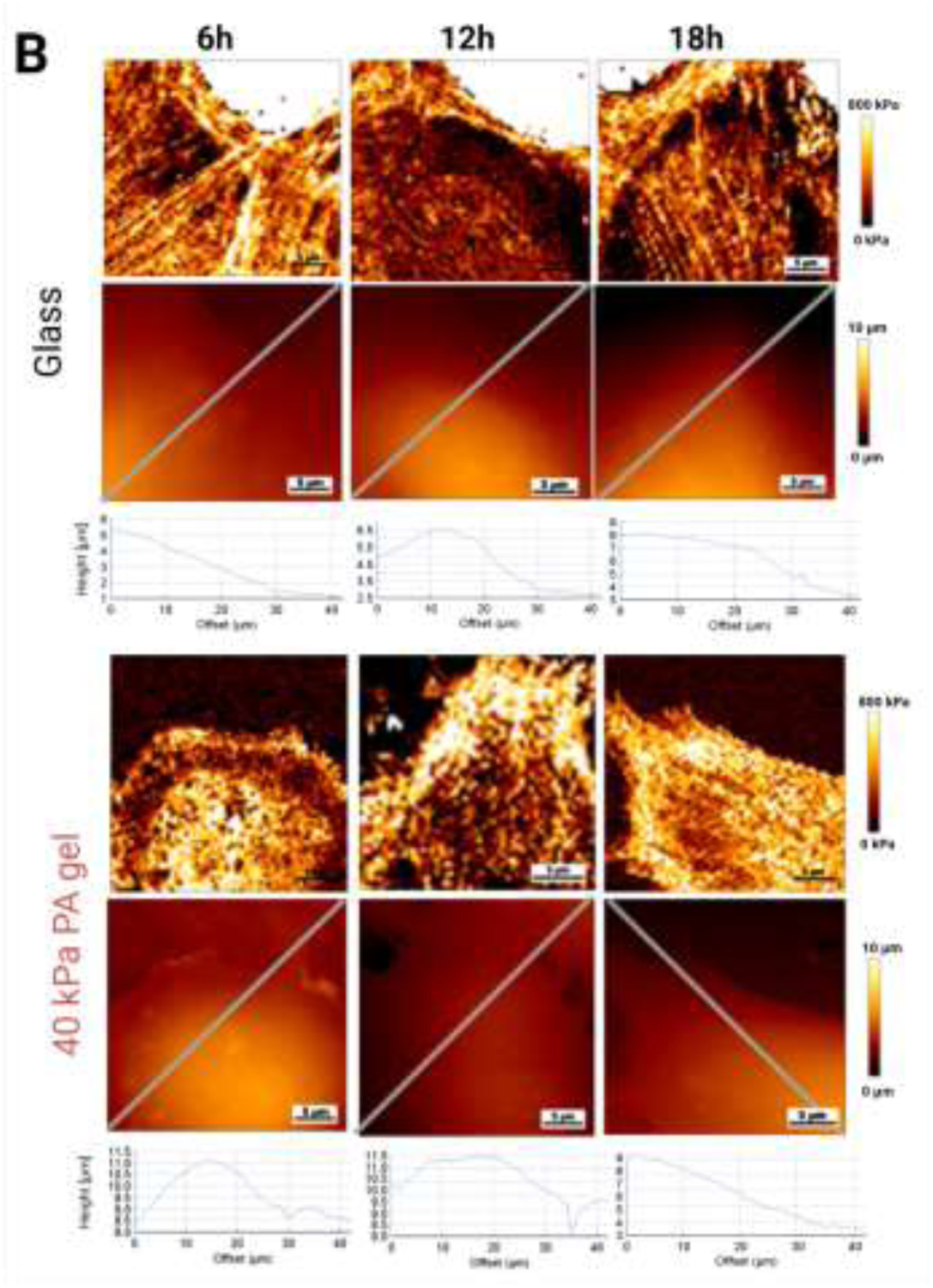
Changes in the elasticity of U2OS cells on substrates of varying stiffness. (A) Box plots showing the elastic modulus (E) of osteosarcoma cells cultured for 6, 12, and 18 h on glass and 40 kPa PA substrates. Measurements were performed on fixed cells. Each point represents the mean E of a single cell. Boxes denote the interquartile range (25th–75th percentile), horizontal lines indicate medians, and squares represent mean values. Statistical significance was determined by two-way ANOVA with Bonferroni post hoc test: *p < 0.01, **p < 0.001; ***p<0.0001. Sample sizes: glass N = 30; 40 kPa PA N = 29. (B) Representative atomic force microscopy (AFM) maps of elastic modulus (E) and height maps and cross-sectional profiles of U2OS cells.

Elasticity maps (Fig. 3B) further corroborated these observations. Cells on glass exhibited prominent, well-defined cytoskeletal fibers (visualized as thick white filaments), reflecting robust cytoskeletal organization. In contrast, cells on 40 kPa PA gels displayed thinner, less intense, and more sparsely distributed cytoskeletal structures. Additionally, substrate stiffness was visually distinguishable in AFM elasticity images: glass regions appeared white, indicative of high stiffness, while 40 kPa substrates showed darker tones (black-orange), corresponding to softer mechanical properties according to the color scale. Height maps and cross-sectional profiles of the cells (Fig. 3B) revealed morphological differences: cells on glass were more flattened, whereas those on 40 kPa gels exhibited increased height and a more rounded morphology, consistent with reduced substrate-induced mechanical tension.

### 3.4. Microparticle Uptake Efficiency Is Modulated by Substrate and Cellular Stiffness

To explore how cells on elastic substate respond to external stimuli, we examined cytoskeletal response to incubation with 2 µm latex beads. U2OS cells were plated on glass-bottom dishes, as well as on 40 kPa PA substrate overnight. Then fluorescent latex microspheres (2 µm) were added and incubated with the cells for 3 hours. Cells were stained for actin and α-tubulin and visualized with confocal microscopy.

U2OS cells exhibit different microtubule-microparticle interaction ways on PA substrates with varying stiffness (Fig. 4A). U2OS cells cultured on glass interacted with the fewest 2 µm microspheres, with most beads remaining at the periphery. In contrast, cells on 40 kPa PA substrate exhibited higher interactions with microspheres, promoting their uptake into the cytoplasm (Fig. 4A). Notably, cells on the 40 kPa PA substrate interacted with the highest number of beads, with an average of 166 microspheres detected within the cell area for 17 cells on images. Quantitative analysis revealed that microtubule-based cup-like structures formed around 6% of beads on glass and 41% on 40 kPa PA gels (Fig. 4B).

**Figure 4.**
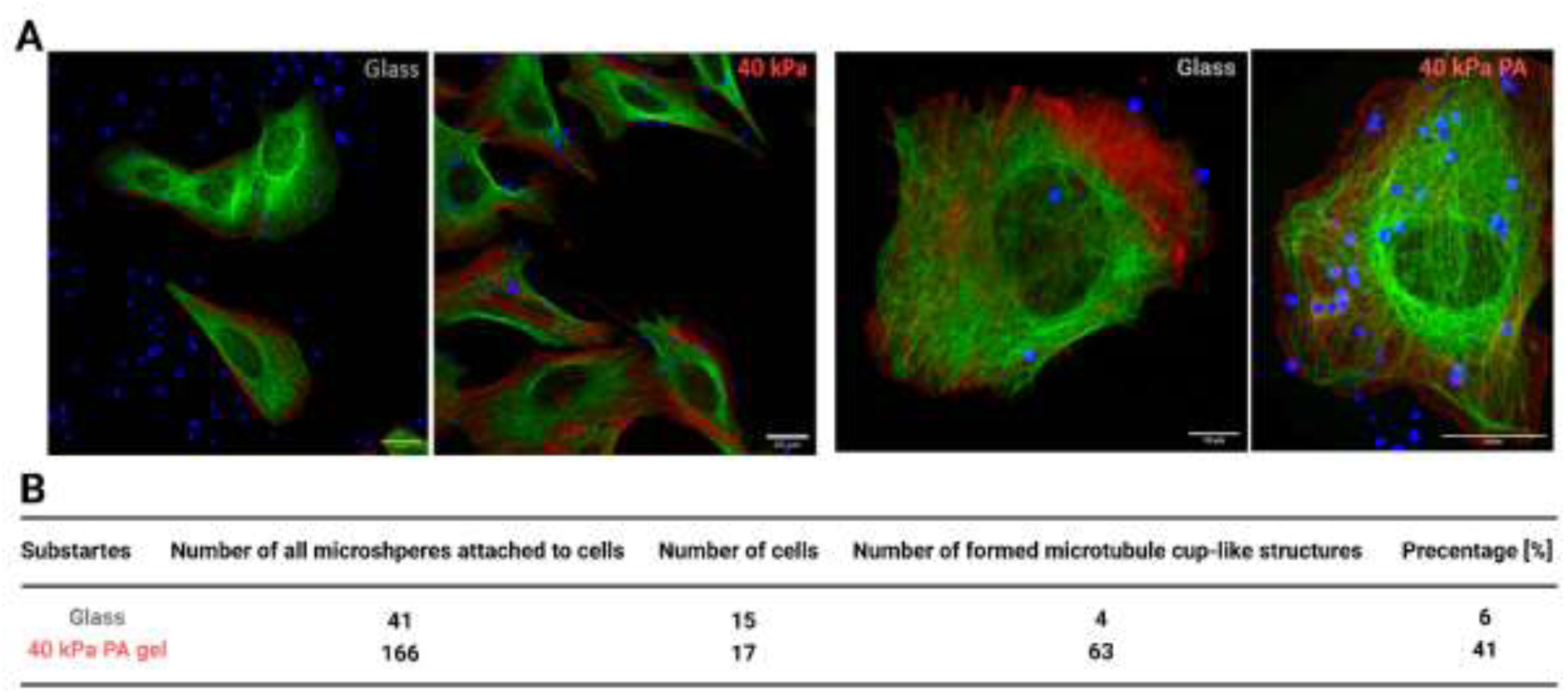
(A) Confocal images showing U2OS cells incubated for 3 h with 2 µm microspheres on glass and 40 kPa PA substrates. Microtubules (green), actin (red), and microspheres (blue) are visualized. Left panels display single-cell views; right panels show larger fields highlighting microsphere distribution and cytoskeletal interactions. (B) Quantification of microtubule-based cup-like structures formed around microspheres across different substrate conditions.

### 3.5. 3D Microtubule Structures Mediate Microspheres Internalization

3D reconstruction analyses of confocal stack images revealed that microtubules in U2OS cells form distinct cup-like structures around 2 µm microspheres. Microtubules were observed surrounding and interacting with microspheres, with orthogonal views confirming their three-dimensional organization (Fig. 5A). The 3D reconstruction further demonstrated the spatial arrangement of these structures, highlighting the close association between microtubules and internalized microspheres (Fig. 5B). 3D views of the reconstruction revealed that microspheres were positioned within the microtubule network rather than simply adhering to the cell surface.

**Figure 5.**
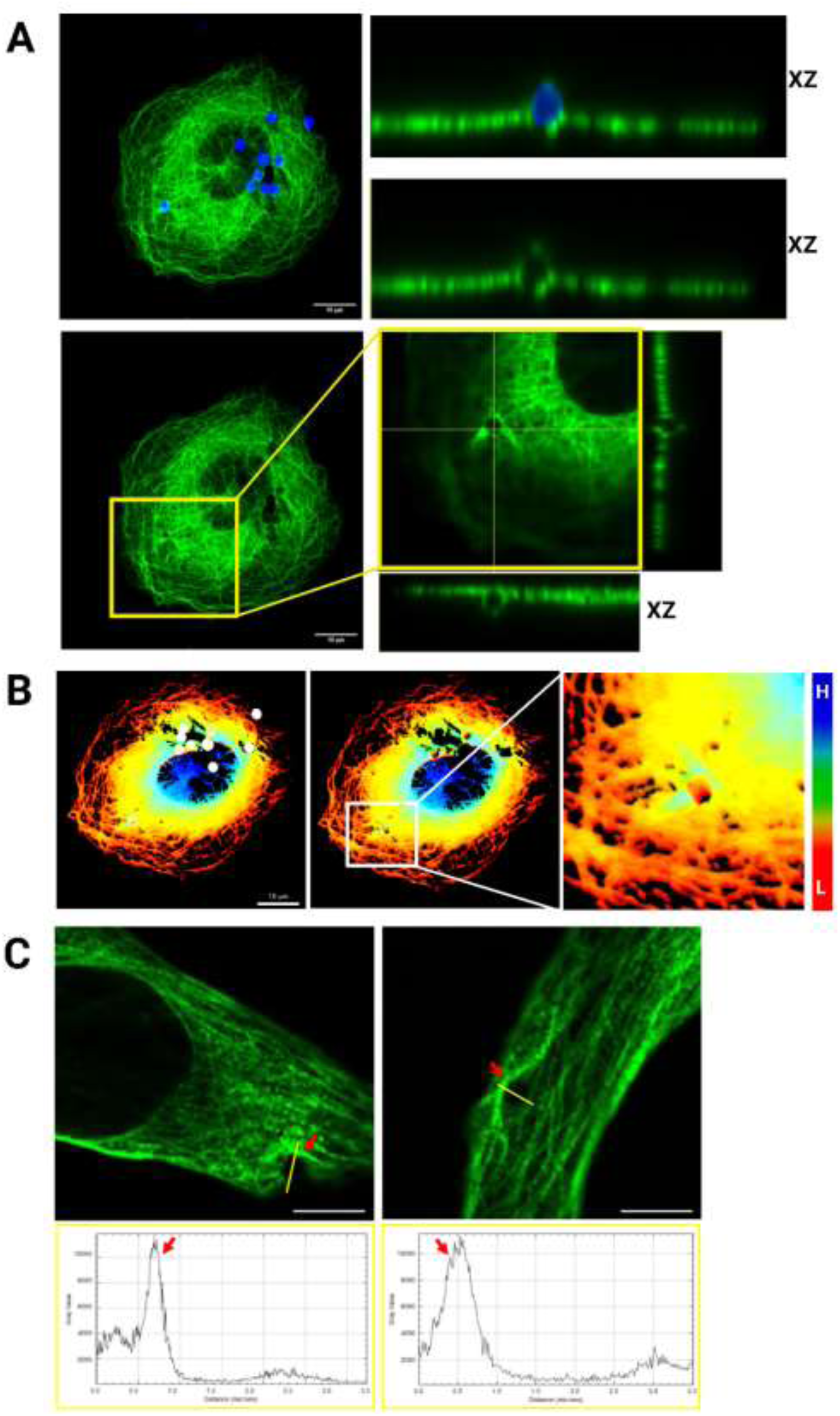
Microtubule-based cup-like structures form around 2 µm microspheres. (A) Representative fluorescence microscopy image of a U2OS cell cultured on a 40 kPa PA gel, stained for microtubules (green) and incubated with 2 µm microspheres (blue) for 3 h. An orthogonal XZ view (right) shows the three-dimensional organization of a microtubule cup-like structure surrounding a microsphere. (B) 3D reconstruction of the microtubule cup structure generated using FluoRender software. The height-color-coding for the same fragment for microtubule filaments (blue → green → yellow → red: High → Low). Rendering parameters are provided in Supplementary file. (C) Representative images highlighting asymmetric formation of the cup-like structure, with elevated regions marked by red arrows. Gray value intensity profiles (pixel brightness) are plotted against distance, illustrating height differences within the structure. Surface plots depict the 3D spatial distribution of microtubule signal intensity. Cells on glass (left) and 40 kPa PA gel (right) are shown.

Additionally, height color-coding analysis (Fig. 5B) showed variations in microtubule organization, with cup-like structures appearing as elevated regions surrounding the beads. A zoomed-in view of the microtubule network further emphasized the localized remodelling of microtubules in response to microsphere interaction. These findings suggest that microtubules actively contribute to particle uptake by forming specialized cup-like structures, supporting their role in endocytosis.

To gain further insights into the spatial organization of these structures, we plotted the z-axis profile and analysed the pixel gray value along this segment. Surface plots were generated specifically for microtubules, providing detailed three-dimensional representations of their distribution and intensity variations. Notably, the microtubule cup-like structures appear to be non-homogeneous. Figure 5C shows that the cup’s side height is heterogeneous, with an elevated cup region being observed. The cup-like structures seem to vary depending on the time of cell-bead interaction or the stiffness of substrate.

### 3.6. Stiffness-Dependent Recruitment and Localization of MAPs

In addition to characterizing microtubule architecture, we investigated the localization of selected microtubule-associated proteins (MAPs) in U2OS cells interacting with 2 µm microparticles on compliant substrates. Specifically, we focused on IQGAP1, MARK2, and CLIP1. Microparticle engagement induced distinct microtubule rearrangements, which in turn influenced MAP localization. While the distribution of MARK2 and CLIP1 remained largely unchanged upon microspheres binding, IQGAP1 was consistently recruited to microtubule-based cup-like structures encasing the beads, suggesting a potential role in their assembly or stabilization. CLIP1 displayed a uniform cytoplasmic distribution throughout the cell, consistent with its established role in global microtubule dynamics. MARK2 predominantly localized to the cytoplasm, with minimal nuclear presence. In contrast, IQGAP1 accumulated at the cell periphery and within discrete cytoplasmic regions, suggesting its potential involvement in localized cytoskeletal remodelling (Fig. 6A).

**Figure 6.**
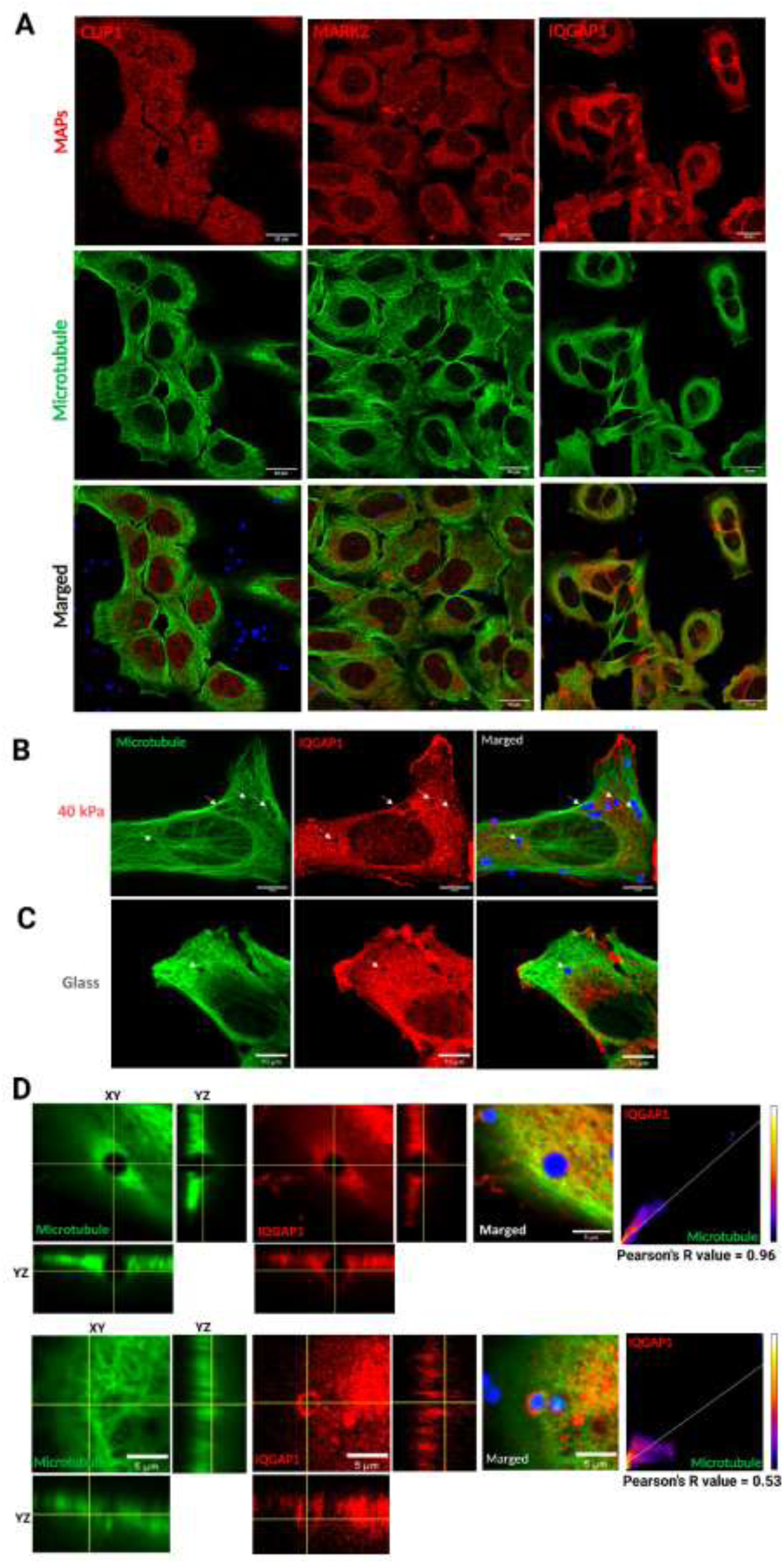
(A) Confocal microscopy images of U2OS cells incubated for 3h with 2 µm microspheres (blue), immunolabeled for microtubules (green) and microtubule-associated proteins (MAPs) — CLIP1, MARK2, and IQGAP1 (red). (B–C) Representative images showing IQGAP1 distribution around microspheres in cells cultured on 40 kPa PA gels (B) and glass substrates (C). Regions of interest, where IQGAP1 colocalizes with microtubules, are indicated by white arrows. (D) Magnified views with orthogonal (XZ) projections illustrate both clear colocalization (upper panel) and spatial dissociation (lower panel) of IQGAP1 from microtubule structures around microspheres. Corresponding Pearson correlation scatter plots demonstrate variability in colocalization efficiency in cells on 40 kPa substrates.

Moreover, we observed that IQGAP1 localized prominently around the 2 µm beads. As shown in Fig. 6B, white arrows indicate distinct regions of IQGAP1 accumulation in U2OS cells cultured on 40 kPa PA gels. Similar but less pronounced patterns were detected on glass substrates (Fig. 6C). Interestingly, in some areas IQGAP1 did not strictly colocalize with the microtubule structures (Fig. 6D), suggesting spatial independence. Notably, IQGAP1 was capable of forming ring-like structures surrounding microspheres even in the apparent absence of microtubules (Fig. 6D, lower panel). Quantitative analysis using Pearson’s correlation coefficient revealed variable colocalization values, ranging from 0.53 to 0.96 in different regions. These findings indicate that IQGAP1 may act independently of microtubules in certain contexts and that its recruitment might be both spatially and temporally regulated. Such ring formation was not observed in cells cultured on glass substrates, although partial colocalization was still evident.

## 4. Discussion

Our study investigates the effects of substrate stiffness on the behavior of U2OS osteosarcoma cells, revealing crucial insights into how mechanical properties (mainly elasticity modulus) of the cellular environment influence cell morphology, cytoskeletal organization, cell elasticity and microspheres uptake.

Osteosarcoma has the potential to metastasize to various organs, including the lungs, bones, and lymph nodes (Shao et al., 2022). These target tissues differ significantly in their mechanical properties, which can be described by their elasticity, typically measured as Young’s modulus. For example, brain tissue is very soft (E = 0.1–1 kPa) (Barnes et al., 2017; Budday et al., 2015), while lung tissue (E = 1–5 kPa), muscle (E = 20–40 kPa) (Basford et al., 2002; Horikawa et al., 1993), and bone (E = 14–23 GPa) (Shoaib et al., 2022) exhibit progressively higher stiffness. Because metastatic osteosarcoma cells may encounter such diverse mechanical environments, it is crucial to model these conditions in vitro. For instance, the medium-soft substrate used in our experiments (E = 38.84 ± 5.59 kPa) closely mimics the elasticity of muscle tissue, a common site of metastasis. In contrast, glass (E = 71.40 ± 11.34 GPa) (Ryou et al., 2020), which is often used as a control surface in cell culture, approximates the stiffness of bone tissue, another frequent target of osteosarcoma dissemination.

### 4.1. Substrate Stiffness Modulates Cell Morphology and Spreading

We observed that U2OS cells cultured on stiff substrates like glass exhibited greater spreading areas over time compared to those on the softer 40 kPa PA substrate. Previous studies have reported that cell spreading, cell morphology, and cytoskeletal organization are affected by substrate stiffness (Chaudhuri et al., 2015). Adherent cells spreading on stiff substrates show increased area due to cytoskeletal fiber assembly and stiffening (Voigt et al., 2023); (Fletcher & Mullins, 2010). The spreading behavior on stiffer substrates can be attributed to the ability of cells to generate higher traction forces, which facilitate a more spread-out morphology (Carotenuto et al., 2022); (Handorf et al., 2015). Yoshikawa H. et al. showed that C2C12 cells adhering to soft or flexible hydrogels can be detached using laser-induced shock waves at lower pressures than cells on rigid hydrogels. This shows that cell adhesion forces are lower on soft substrates, so cells can be easily detached (Yoshikawa et al., 2011). These findings suggest that stiffer substrates promote greater and more sustained spreading of U2OS cells.

### 4.2. Cytoskeletal Architecture Is Altered by Substrate Stiffness

Confocal microscopy analysis revealed that actin stress fibers and overall cytoskeletal architecture were more pronounced in U2OS cells cultured on stiffer substrates. These findings are in line with studies showing that cytoskeletal assembly and organization scale with substrate stiffness (Solon et al., 2007; Voigt et al., 2023). Furthermore, confocal microscopy images showed that osteosarcoma cells exhibit large and well-organized actin stress fibers on 40 kPa PA and glass (Fig.2). Previous studies suggest that fibroblasts on ECM functionalized compliant PA gels (<5 kPa) show mostly cortical actin, but no actin stress fibers. Actin stress fibers become visible on stiff PA gels (>10 kPa), well organized stress fiber bundles are apparent on rigid glass substrate in a stiffness dependent manner (Solon et al., 2007). As actin and microtubule networks are crucial for transmitting mechanical signals and maintaining structural integrity, such remodelling likely reflects cellular adaptation to increased mechanical demands (Fletcher & Mullins, 2010; Yi et al., 2022).

### 4.3. Substrate Stiffness Influences Cell Elasticity

AFM measurements demonstrated that the elastic modulus of U2OS cells was higher on glass than 40 kPa PA substrate, indicating enhanced cellular stiffness. This is consistent with increased cytoskeletal organization and spreading, which are known to correlate with mechanical robustness and functional capacity (Lekka, 2016). The larger amplitude of stiffness fluctuations on glass may be due to the dynamic remodeling of the actin cortex under higher mechanical loads. Our findings mirror prior observations in osteosarcoma cells, where survival and stemness were favoured on stiffer substrates (Mylona et al., 2008; Jabbari et al., 2015). These stiffness-dependent mechanical profiles could influence migration, metastasis, and resistance to therapy.

### 4.4. Microparticle Uptake Efficiency Is Modulated by Substrate and Cellular Stiffness

We found that U2OS cells cultured on 40 kPa substrates exhibited the highest interaction with 2 µm microspheres (166 beads per 17 cells), compared to glass (41 beads per 15 cells). On glass, beads predominantly remained at the cell periphery, whereas cells on 40 kPa PA more efficiently internalized them into the cytoplasm. Glass substrates may reduce microspheres internalization by U2OS due to excessive cortical tension or overly rigid actin and microtubules architecture, which impairs protrusions formation and uptake. These results align with prior studies demonstrating increased particle uptake on softer substrates due to enhanced membrane flexibility and cytoskeletal plasticity (Lee et al., 2022; Visonà et al., 2024).

### 4.5. 3D Microtubule Structures Mediate Microspheres Internalization

A key finding of our study is the identification of microtubule-based, cup-like structures forming around microspheres—particularly prominent on softer substrates (41% on 40 kPa PA vs. 6% on glass). This may represent the first documentation of such microtubule-based engulfment structures in U2OS cells. These structures resemble phagocytic cups and suggest an remarkable, active role for microtubules in particle internalization in U2OS cells. This observation extends findings from Adamczyk et al. (2021), who reported similar structures in fibroblasts, and adds new evidence that microtubules, not just actin filaments, can participate directly in endocytic processes. Unlike traditional views of microtubules as intracellular transport tracks (Matis, 2020), our data suggest a cytoskeletal adaptation mechanism involving surface remodelling. In some cases, asymmetry in cup elevation suggests localized directional forces, potentially arising from intracellular tension, time-dependency or asymmetric bead binding.

### 4.6. Stiffness-Dependent Recruitment and Localization of MAPs

IQGAP1 is a well-characterized scaffold protein that connects actin filaments and microtubules, playing key roles in migration, adhesion, and signaling (Sivaraj et al., 2022; Ren et al., 2007). Notably, IQGAP1 was also observed forming rings around microspheres even in the absence of microtubules, indicating that actin-based mechanisms may also mediate its localization. These observations are supported by earlier reports: IQGAP1 localizes to actin coats on vesicles (Müller et al., 2019) and accumulates in microtubule-depleted zones or ER-excluded regions (Samson et al., 2017). The fact that IQGAP1 is overexpressed in multiple cancers (Hedman et al., 2015) suggests its mechanosensitive role could contribute to tumor invasiveness and might be a potential therapeutic target.

Additionally, microtubule-associated proteins (MAPs) such as MARK2 and CLIP1 are known regulators of microtubule dynamics. MARK2 (microtubule affinity-regulating kinase 2) phosphorylates microtubule-associated proteins and modulates microtubule stability and orientation, influencing cell polarity and directional transport (Conboy et al., 2024). CLIP1 (also known as CLIP-170) binds to the plus ends of growing microtubules and facilitates their interaction with other cellular structures, playing a critical role in microtubule tracking and stabilization (Lansbergen & Akhmanova, 2006). Although neither protein exhibited targeted enrichment around microtubule cups in our study, their widespread cytoplasmic distribution may support general microtubule regulation during mechanosensitive remodeling.

## 5. Conclusions

Our study demonstrates that substrate stiffness, especially at an intermediate level of 40 kPa, has a profound effect on osteosarcoma (U2OS) cell morphology, elasticity, and cytoskeletal organization. Cells cultured on this stiffness displayed greater adaptability and internalization capacity for 2 µm microspheres, suggesting enhanced cytoskeletal plasticity. We also identified the formation of novel microtubule cup-like structures during particle uptake, with frequent localization of the microtubule-associated protein IQGAP1, pointing to its active role in mechanosensitive cytoskeletal remodelling. These findings underscore how finely tuned mechanical environments influence cancer cell behaviour and offer valuable insights into mechanotransduction processes relevant for cancer progression and the design of functional biomaterials.

## Author Contributions

CR designed and performed all experiments, analyzed data, and wrote the manuscript. AK and MTK performed AFM experiments, acquisition of data, and analysis and interpretation of data. DP analyzed microscopic images and data. ZR and MTK devised and supervised this study. DP, AK, ZR and MTK critically read and revised the manuscript. All authors have accepted responsibility for the entire content of this manuscript and approved its submission.

## Acknowledgments

This research was founded by the “Research support module” as part of the “Excellence Initiative – Research University” program at the Jagiellonian University in Kraków no. RSM/63/RC (to RC).

## Abbreviations

AFM: Atomic force microscopy
FBS: Fetal bovine serum
MAPs: Microtubule-associated proteins
OS: Osteosarcoma
PA: Polyacrylamide
PBS: Phosphate-buffered saline

