## Supplementary materials for "Substrate elasticity regulates cytoskeletal remodeling and mechanical behavior of U2OS osteosarcoma cells"

1. The elasticity of a 40 kPa polyacrylamide gel substrate.

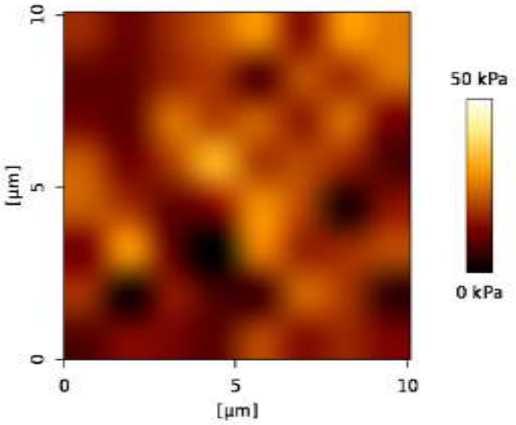


Figure S1. Confirmation of Young Modulus of PA gel. Topography AFM map of 40 kPa PA gels.

2. Time-lapses of the live cells 40kPa (Video 01), and glass (Video 02) were taken using Wide-field microscopy Zeiss Axio Observer Z1 (Carl Zeiss).
3. Statistical analysis.

Tab.S1. Quantification of adherent cell area demonstrating the difference in spreading area between cells on 40 kPa gel and glass substrate. Only significant differences are shown in the table.

| Substrate | Time | vs | Substrate | Time | Mean | SEM | t Value | Probability | Significant |
|---|---|---|---|---|---|---|---|---|---|
| glass | 6h | vs | glass | 0h | 576,40168 | 83,3237 | 6,91762 | <0.0001 | 1 |
| glass | 12h | vs | glass | 0h | 935,61197 | 89,82535 | 10,4159 | <0.0001 | 1 |
| glass | 12h | vs | glass | 6h | 359,21029 | 66,33761 | 5,41488 | <0.0001 | 1 |
| glass | 18h | vs | glass | 0h | 1084,8503 | 90,49162 | 11,98841 | <0.0001 | 1 |
| glass | 18h | vs | glass | 6h | 508,44862 | 67,23703 | 7,56203 | <0.0001 | 1 |
| glass | 24h | vs | glass | 0h | 1063,21793 | 91,21291 | 11,65644 | <0.0001 | 1 |

| | | | | | | | | | |
|---|---|---|---|---|---|---|---|---|---|
| glass | 24h | vs | glass | 6h | 486,81625 | 68,20469 | 7,13758 | <0.0001 | 1 |
| 40kPa PA | 0h | vs | glass | 6h | -523,7967 | 71,60895 | -7,31468 | <0.0001 | 1 |
| 40kPa PA | 0h | vs | glass | 12h | -883,00699 | 79,07968 | -11,16604 | <0.0001 | 1 |
| 40kPa PA | 0h | vs | glass | 18h | -1032,24532 | 79,83568 | -12,92962 | <0.0001 | 1 |
| 40kPa PA | 0h | vs | glass | 24h | -1010,61295 | 80,65233 | -12,53049 | <0.0001 | 1 |
| 40kPa PA | 6h | vs | glass | 0h | 522,00494 | 90,49162 | 5,76854 | <0.0001 | 1 |
| 40kPa PA | 6h | vs | glass | 12h | -413,60703 | 75,14369 | -5,50422 | <0.0001 | 1 |
| 40kPa PA | 6h | vs | glass | 18h | -562,84536 | 75,93888 | -7,41182 | <0.0001 | 1 |
| 40kPa PA | 6h | vs | glass | 24h | -541,21299 | 76,79697 | -7,04732 | <0.0001 | 1 |
| 40kPa PA | 6h | vs | 40kPa PA | 0h | 469,39996 | 79,83568 | 5,87958 | <0.0001 | 1 |
| 40kPa PA | 12h | vs | glass | 0h | 679,25908 | 90,84497 | 7,47712 | <0.0001 | 1 |
| 40kPa PA | 12h | vs | glass | 18h | -405,59122 | 76,3596 | -5,31159 | <0.0001 | 1 |
| 40kPa PA | 12h | vs | glass | 24h | -383,95885 | 77,21302 | -4,97272 | <0.0001 | 1 |
| 40kPa PA | 12h | vs | 40kPa PA | 0h | 626,6541 | 80,23597 | 7,81014 | <0.0001 | 1 |
| 40kPa PA | 18h | vs | glass | 0h | 874,82397 | 92,85061 | 9,42184 | <0.0001 | 1 |
| 40kPa PA | 18h | vs | glass | 6h | 298,42229 | 70,37984 | 4,24017 | 0,00272 | 1 |
| 40kPa PA | 18h | vs | 40kPa PA | 0h | 822,21899 | 82,49994 | 9,9663 | <0.0001 | 1 |
| 40kPa PA | 18h | vs | 40kPa PA | 6h | 352,81903 | 78,73511 | 4,48109 | 9,36539E-4 | 1 |
| 40kPa PA | 24h | vs | glass | 0h | 865,54452 | 93,78569 | 9,22896 | <0.0001 | 1 |
| 40kPa PA | 24h | vs | glass | 6h | 289,14284 | 71,60895 | 4,0378 | 0,00641 | 1 |
| 40kPa PA | 24h | vs | 40kPa PA | 0h | 812,93954 | 83,55093 | 9,72987 | <0.0001 | 1 |
| 40kPa PA | 24h | vs | 40kPa PA | 6h | 343,53957 | 79,83568 | 4,30308 | 0,00207 | 1 |

Tab.S2. Quantification of AFM measurements. Elastic modulus of cells on 40 kPa gel and glass substrate.

| **Substrate** | **Time** | **vs.** | **Substrate** | **Time** | **Mean** | **SEM** | **t Value** | **Probability** | **Significant** |
|---|---|---|---|---|---|---|---|---|---|
| glass | 12h | vs. | glass | 6h | 70,84862 | 6,61001 | 2,87E-04 | 0,05 | 1 |
| glass | 18h | vs. | glass | 6h | 72,88119 | 2,27443 | 0,59688 | 0,05 | 0 |
| glass | 18h | vs. | glass | 12h | 74,50298 | 4,06088 | 0,06161 | 0,05 | 0 |
| 40kPa PA | 6h | vs. | glass | 6h | 72,88119 | 7,33681 | <0.0001 | 0,05 | 1 |
| 40kPa PA | 6h | vs. | glass | 12h | 74,50298 | 0,89131 | 0,98823 | 0,05 | 0 |
| 40kPa PA | 6h | vs. | glass | 18h | 76,43844 | 4,82679 | 0,01487 | 0,05 | 1 |
| 40kPa PA | 12h | vs. | glass | 6h | 70,84862 | 8,44444 | <0.0001 | 0,05 | 1 |
| 40kPa PA | 12h | vs. | glass | 12h | 72,51587 | 1,79225 | 0,80113 | 0,05 | 0 |
| 40kPa PA | 12h | vs. | glass | 18h | 74,50298 | 5,80532 | 0,00186 | 0,05 | 1 |
| 40kPa PA | 12h | vs. | 40kPa PA | 6h | 74,50298 | 0,85313 | 0,99035 | 0,05 | 0 |
| 40kPa PA | 18h | vs. | glass | 6h | 70,84862 | 4,96922 | 0,01117 | 0,05 | 1 |
| 40kPa PA | 18h | vs. | glass | 12h | 72,51587 | 1,60307 | 0,86515 | 0,05 | 0 |
| 40kPa PA | 18h | vs. | glass | 18h | 74,50298 | 2,50056 | 0,49451 | 0,05 | 0 |
| 40kPa PA | 18h | vs. | 40kPa PA | 6h | 74,50298 | 2,45163 | 0,51645 | 0,05 | 0 |
| 40kPa PA | 18h | vs. | 40kPa PA | 12h | 72,51587 | 3,39532 | 0,17458 | 0,05 | 0 |

4. Cup-like structures of actin microtubules.

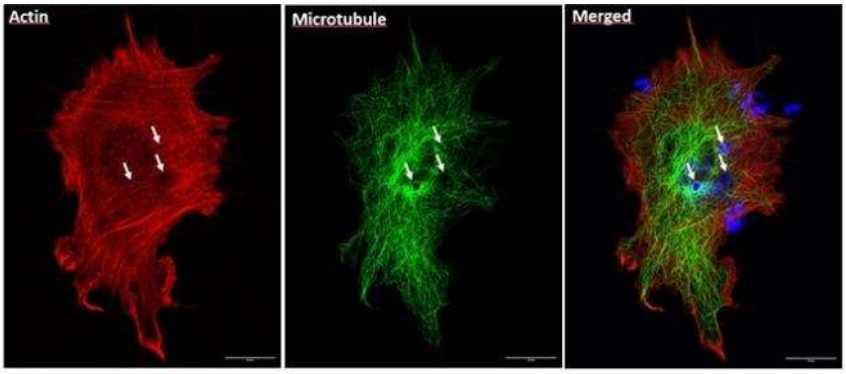


Figure S2. Exemplary images showing significant cytoskeletal remodeling occurred within 3 hours, including the formation of distinct "holes" in the network involving both actin filaments (red) and microtubules (green). The most visible differences are highlighted with white arrows.

5. 3D rendering parameters.

Table S3. FluoRender software analysis settings.

| **Florender Settings** | Microtubules | Microparticles |
|---|---|---|
| Gamma | 0.14 | 0.33 |
| Saturation | 4947 | 17953 |
| Luminescence | 21724 | 17241 |
| Alpha | 17953 | 17953 |
| Extract Boundary | 0.0025 | 0.0555 |
| Threshold | 5827 - 20609 | 0 - 20609 |
| Sample Rate | 2.0 | 2.0 |
| Colormap | 1400 - 20700 | - |

6. MAPs proteins localization.

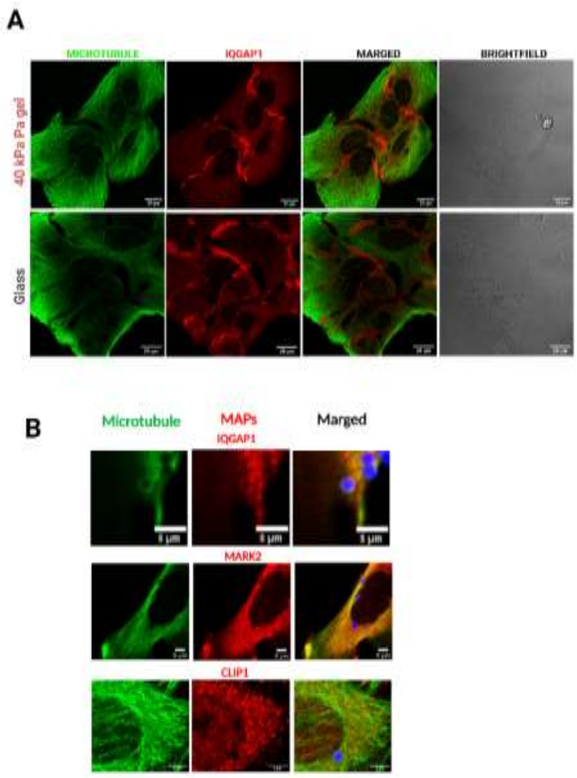


Figure S3. Distribution of MAPs in U2OS cells. (A) Control U2OS cells cultured on glass and immunostained for microtubules and IQGAP1, without microsphere incubation. (B) High-magnification images of individual cells incubated with microspheres, showing the distribution of MAPs, including IQGAP1, MARK2, and CLIP1.

7. Cells viability.

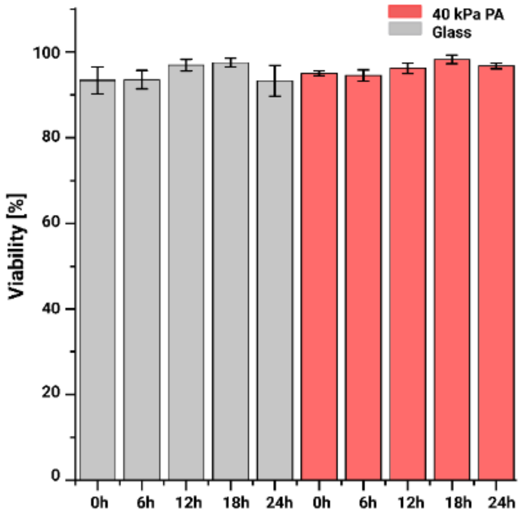


Figure S4. Cell viability on substrates of different stiffness. Bar graph showing the percentage of viable U2OS cells cultured for 24 h on glass and 40 kPa polyacrylamide (PA) gel. Viability was assessed using the LIVE/DEAD™ Viability/Cytotoxicity Assay Kit. Cell viability remained high (>95%) on both substrates.
